# Analysis of RNA processing directly from spatial transcriptomics data reveals previously unknown regulation

**DOI:** 10.1101/2023.03.13.532412

**Authors:** Julia Olivieri, Julia Salzman

## Abstract

Technical advances have led to an explosion in the amount of biological data available in recent years, especially in the field of RNA sequencing. Specifically, spatial transcriptomics (ST) datasets, which allow each RNA molecule to be mapped to the 2D location it originated from within a tissue, have become readily available. Due to computational challenges, ST data has rarely been used to study RNA processing such as splicing or differential UTR usage. We apply the ReadZS and the SpliZ, methods developed to analyze RNA process in scRNA-seq data, to analyze spatial localization of RNA processing directly from ST data for the first time. Using Moran’s I metric for spatial autocorrelation, we identify genes with spatially regulated RNA processing in the mouse brain and kidney, re-discovering known spatial regulation in *Myl6* and identifying previously-unknown spatial regulation in genes such as *Rps24*, *Gng13*, *Slc8a1*, *Gpm6a*, *Gpx3*, *ActB*, *Rps8*, and *S100A9*. The rich set of discoveries made here from commonly used reference datasets provides a small taste of what can be learned by applying this technique more broadly to the large quantity of Visium data currently being created.

## Introduction

One of the most fundamental questions of RNA biology is “where can I find this RNA in an organism?” On a large scale, this question can be answered by sequencing different tissues of an organism and characterizing their RNA compositions. However, a more fine-grained answer was out of reach until the introduction of single-cell RNA sequencing (scRNA-seq) (Tang et al., 2009) and spatial transcriptomics (ST) (Ståhl et al., 2016), each of which provide complementary but distinct answers. In scRNA-seq, individual RNAs are labeled with their cell of origin, but the relative spatial locations of these cells are unknown. In ST, RNAs are labeled with cartesian coordinates describing their spatial locations, but the resolution is not yet able to distinguish individual cells. While scRNA-seq requires time-intensive cell type annotation before most analysis, ST data allows annotation-free analysis immediately. The promise of ST methods led to spatially resolved transcriptomics being named the “Method of the Year” by Nature in 2020 (Marx, 2021).

Differential RNA processing, including alternative splicing and differential UTR usage, is ubiquitous (Wang et al., 2008), highly regulated (Olivieri et al., 2021), and implicated in human disease (Anczuków & Krainer, 2016) and treatment (Mendell et al., 2017). Understanding the spatial localization of RNA isoforms would help disentangle effects resulting from RNA processing from those resulting in gene expression difference (Figure 1A). Despite its importance, the function of the vast majority of alternative RNA processing is not understood (Wan & Larson, 2018). Despite rising interest in ST, analysis of RNA processing in ST data is almost completely unexplored. ST data suffers from challenges related to sparsity and the 3’ bias of the reads that impeded RNA processing analysis in droplet-based scRNA-seq data (Arzalluz-Luque & Conesa, 2018). When RNA splicing has been studied in ST data, results have relied on integration with separate long-read or full-length-coverage data that is then mapped to spatial locations algorithmically (Booeshaghi et al., 2021; Joglekar et al., 2021; Lebrigand et al., 2020). These methods each require a matched dataset for RNA processing analysis in ST data, which makes them inapplicable to most of the ST datasets publicly available. There has been some analysis of 3’ UTR localization in subcellular spatial data (Bierman & Salzman, 2022), but most relies on seqFISH experiments rather than ST data (Cassella & Ephrussi, 2022). Up to this point, analysis of RNA processing directly from ST data has been out of reach.

**Figure 1.**
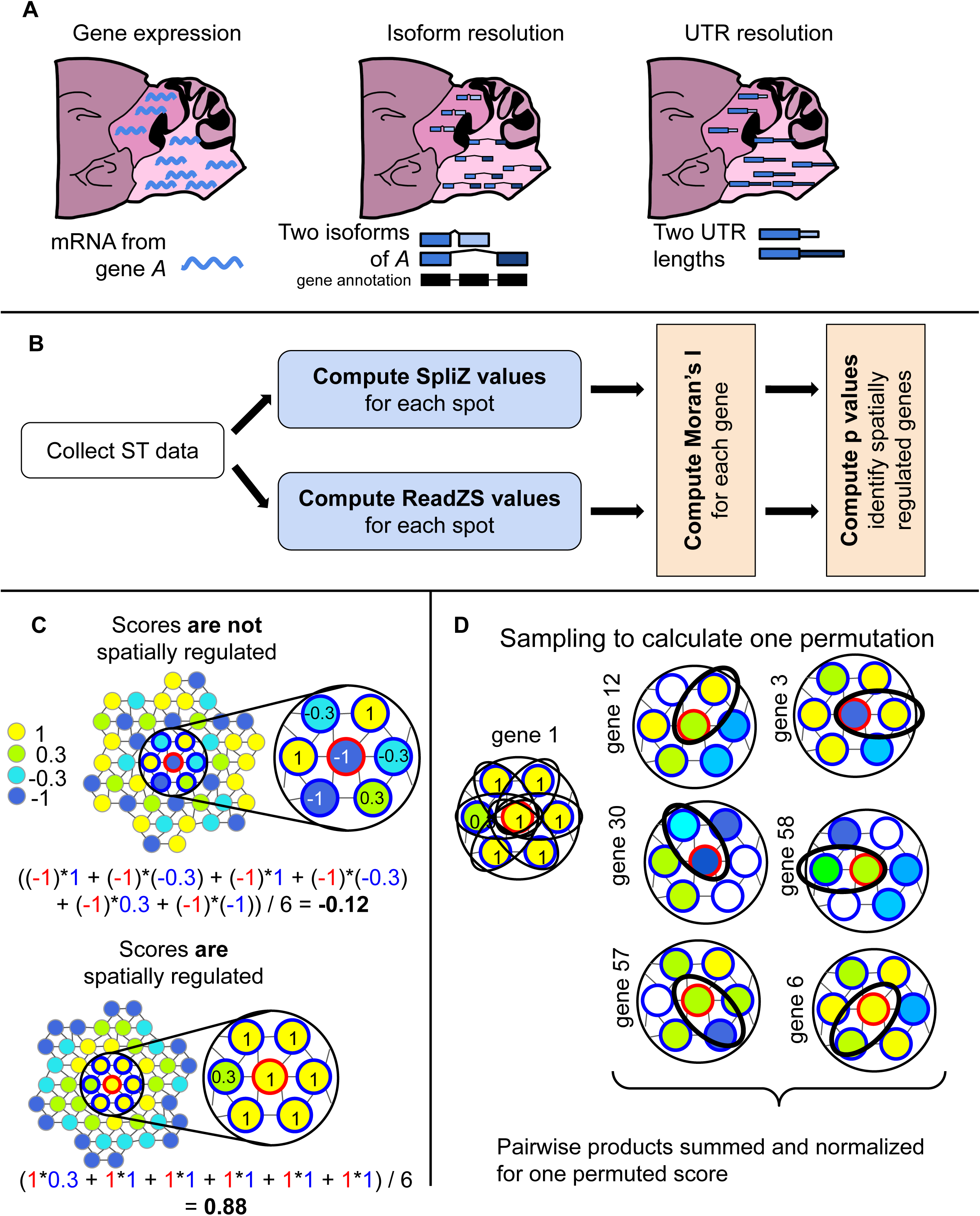
A) Analysis of ST data has almost exclusively focused on finding spatial regulation of gene expression, missing potential spatial regulation of RNA processing and splicing. B) Starting with ST data, our pipeline calculates SpliZ and ReadZS values independently for all spots, and then determines statistical significance using Moran’s I with permutations. C) An example calculation using Moran’s i for all neighbors (outlined in blue) of the spot outlined in red. The red spot’s score is multiplied with each blue spot’s score individually, and the results are summed and normalized by the number of pairs. The pattern on the left results in a score close to zero, while the pattern on the right results in a score close to one. This indicates that the pattern on the right is more likely to have spots with similar scores grouped together. For the full metric, this calculation is performed over all neighboring pairs (rather than only for the neighbors of one spot). D) To calculate p values, permutations are performed by sampling the scores of each neighboring pair from the scores of that pair in a random gene.

In this work we apply the SpliZ (Olivieri et al., 2022) and ReadZS (Meyer et al., 2022) methods, developed to analyze alternative splicing and 3’ UTR usage, in scRNA-seq data, to study spatial regulation of RNA processing directly from ST data for the first time (Figure 1B). Because both droplet-based scRNA-seq and ST technologies suffer from similar challenges due to sparsity and 3’ bias, these methods are directly applicable to ST data. We use Moran’s I measure of spatial autocorrelation with a carefully constructed null to identify genes with spatially regulated RNA processing in the mouse brain and kidney.

## Results

We applied the SpliZ and ReadZS, recently-developed methods to identify splicing differences and UTR length differences in single-cell data respectively (Meyer et al., 2022; Olivieri et al., 2022), to identify spatially-regulated RNA processing changes in the mouse brain and kidney. Although these methods were not developed specifically for application to ST data, none of the specificities of spatial transcriptomics data invalidate the assumptions of the SpliZ or ReadZS: they are already robust to low sampling rates and unknown capture biases (Meyer et al., 2022; Olivieri et al., 2022). We developed new methods to identify spatial regulation of these scores to take into account the spatial coordinates of the data and potential mixing between neighboring spots (Methods). Intuitively, the ReadZS quantifies the average location of read build-up in discrete genomic windows (5000 bp-length continuous regions of the genome) in each spatial cell, while the SpliZ quantifies the deviation of the ranked length of introns in a given cell from the population average for that gene (lower values indicate shorter introns, higher values indicate longer introns). ST data has similar formatting and biases compared to droplet-based scRNA-seq data, meaning that we can apply the ReadZS and SpliZ to ST data without modification, though neither has been applied to spatial data to this point.

The ReadZS and SpliZ were chosen for this analysis because both of these methods provide an individual score for each cell-gene pair, which is easily adapted to providing a score for each spot-gene pair. Both of these methods take into account reads across the gene body, though this is limited by the 3’ bias of the data. Due to the sparsity and 3’ bias of Visium data, approaches designed to analyze RNA processing in full-length sequencing analysis are not applicable. The SpliZ and ReadZS are two of the limited number of tools available that are designed for the analysis of RNA processing in droplet-based data. Other available tools tend to rely on aggregating data across multiple cells using a method called pseudo-bulking (W. V. Li et al., 2021; Patrick et al., 2020). Pseudo-bulking methods can help increase power and detect changes for genes with very low read counts, but at the cost of potentially obscuring subtle spatial patterns in the data. Others are based on PSI measurements, which are vulnerable to artifacts due to sparsity (Buen Abad Najar et al., 2020; Olivieri et al., 2022; Wen et al., 2022).

We applied the SpliZ and ReadZS to Visium data from the sagittal-posterior mouse brain (2 biological replicates), the sagittal-anterior mouse brain (2 biological replicates), and the mouse kidney (1 biological replicate) (Methods). The number of spots covered by the tissue in these datasets ranges from 1,436 to 3,355, and sequencing depth is generally lower than for 10x data, leading to fewer genes and genomic windows with ReadZS and SpliZ scores per spot (Table 1) (Olivieri et al., 2021).

**Table 1:** Dataset summary. Columns are: `dataname` (name of dataset), `num_spots` (the number of Visium spots with tissue information), `med_reads_per_spot` (median number of reads per spot), `SpliZ_med_per_spot` (median number of genes with SpliZ values per spot), `ReadZS_med_per_spot` (median number of 5000-bp windows with ReadZS values per spot), `ge_med_per_spot` (median number of genes with nonnegative gene expression per spot), and `ReadZS_ge_med_per_spot` (median number of 5000-bp windows with nonnegative gene expression per spot)

Significantly spatially regulated RNA processing can be quantified using Moran’s I measure of spatial autocorrelation (Getis, 2007) on SpliZ and ReadZS scores. Applying the SpliZ and ReadZS to ST data results in a single ReadZS score per genomic window per spot, and a single SpliZ score per gene per spot (Methods). For each gene, Moran’s I quantifies the average similarity between the value of each neighboring pair of spots (Methods). A negative value implies neighboring pairs are more likely to have different scores and a positive value implies neighboring pairs are more likely to have similar scores. This paper focuses on genes with high Moran’s I values (Figure 1C). Significance values were calculated through permutation testing, with modifications to avoid high type I error rates (Figure 1D, Methods, Chung & Romano, 2013). Genes with significant Moran’s I scores (p value < 0.05 by permutation testing, Methods) in both biological replicates have highly correlated Moran’s I scores for the SpliZ (Spearman correlation 0.936-0.943) and the ReadZS (Spearman correlation 0.765-0.885), indicating that this metric is robust and replicable (Figure 2—figure supplement 1, Methods).

**Figure 2.**
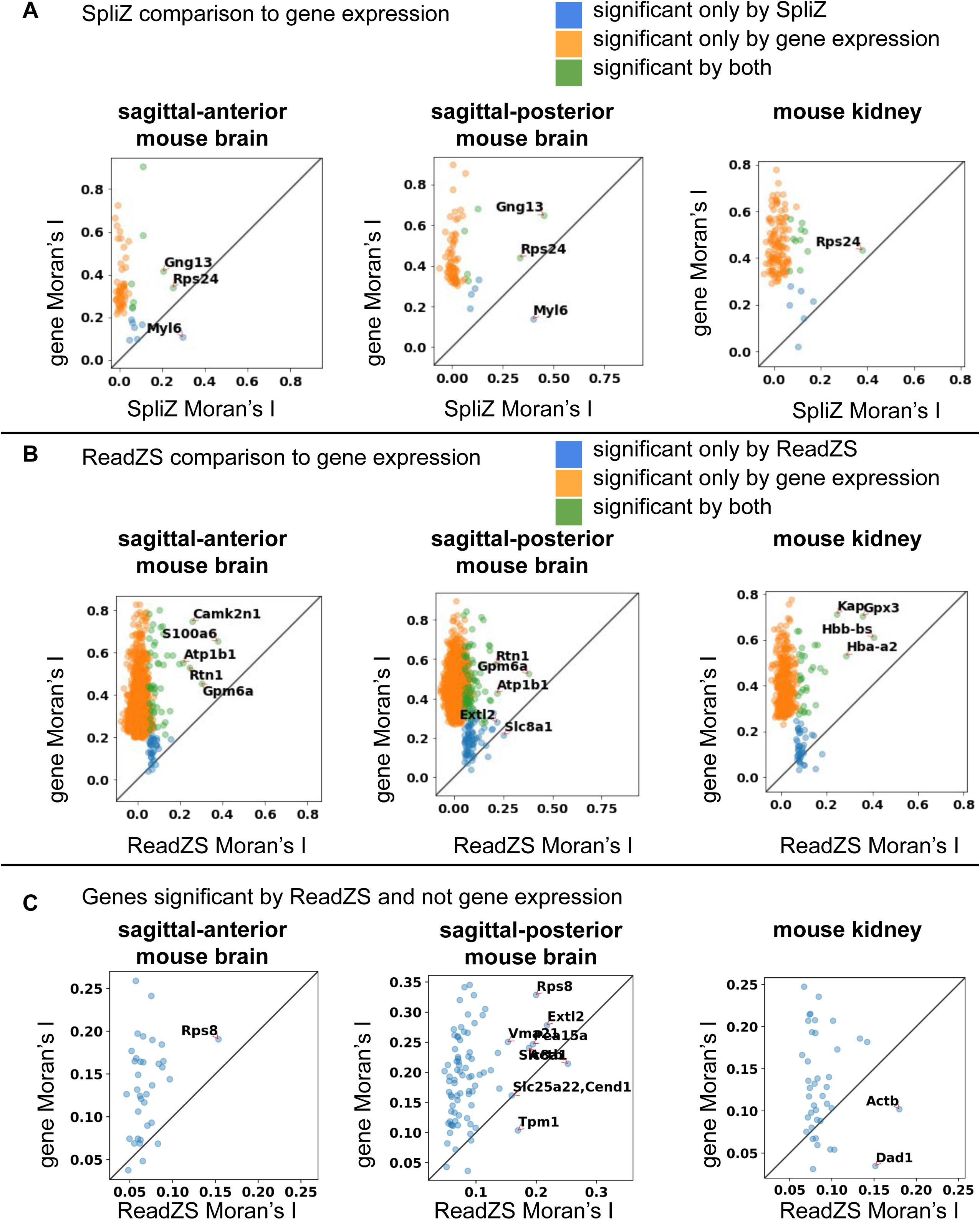
A) For each mouse dataset, Moran’s I for the SpliZ is included on the x axis and Moran’s I for gene expression is included on the y axis. Points with Moran’s I for the SpliZ > 0.2 are labeled. In several genes (marked in blue), Moran’s I for the SpliZ is larger than for gene expression, indicating that the spatial regulation of splicing is more pronounced than the spatial regulation of gene expression. B) Same as A, except Moran’s I for the ReadZS is included on the x axis. C) These plots show only the points from B) corresponding to genomic windows that are significant by the ReadZS but not gene expression. Points for which Moran’s I for ReadZS > 0.15 are labeled. Note that the difference in thresholds in this figure only correspond to differences in which gene names are included in the plot for visual clarity, and do not affect the rest of the analysis.

Despite the 3’ bias and sparsity of ST data, applying Moran’s I to sections of the mouse brain revealed that splicing in 0.8-2.2% of detected genes (8-17 genes) and RNA processing in 1.1-5.5% of detected genomic windows (57-161 windows) are significantly spatially regulated (Table 2). Although this is lower than the 13-22% of detected genes for which the gene expression was spatially regulated, it still shows that spatial regulation of RNA processing can be analyzed in ST data. In each tissue section the SpliZ and ReadZS identified genes for which RNA processing was significantly spatially regulated, but gene expression was not (Figure 2; Figure 2—figure supplement 2). Note that fewer reads are used to compute the SpliZ and ReadZS compared to gene expression, which could account for some of this difference.

**Table 2:** Summary of discoveries. Columns are: `dataname` (name of dataset), `score` (either `ReadZS_norm`, `SpliZ_norm`, `ge_norm`, which is gene expression, or `ReadZS_ge_norm`, which is 5000-bp-window expression), `num_genes` (the number of genes tested, or windows in the case of ReadZS_norm or ReadZS_ge_norm), `num_sig` (the number with significant spatial patterns by Moran’s I), `top` (the top 10 genes/windows found to be significant in the dataset for the given score), and `frac_sig` (the fraction of significant genes or windows).

The genes *Myl6*, *Rps24*, and *Gng13* consistently had the highest Moran’s I scores for the SpliZ in both the sagittal-anterior and sagittal-posterior mouse brain (Figure 2A, Table 3). *Myl6* was the only gene that consistently had a higher splicing Moran’s I than gene expression Moran’s I. *Myl6* is a myosin light chain gene that has a known exon skipping event which is cell-type-specifically regulated in human, mouse, and mouse lemur (Olivieri et al., 2021), and was previously reported to be spatially localized in the mouse brain (Lebrigand et al., 2020). The exon-included isoform is most highly expressed in the brainstem and cerebellum, while the exon-excluded isoform is most highly expressed in the cerebrum. Gene expression is more uniform across the brain (Figure 3A; Figure 3—figure supplements 1 and 2).

**Figure 3.**
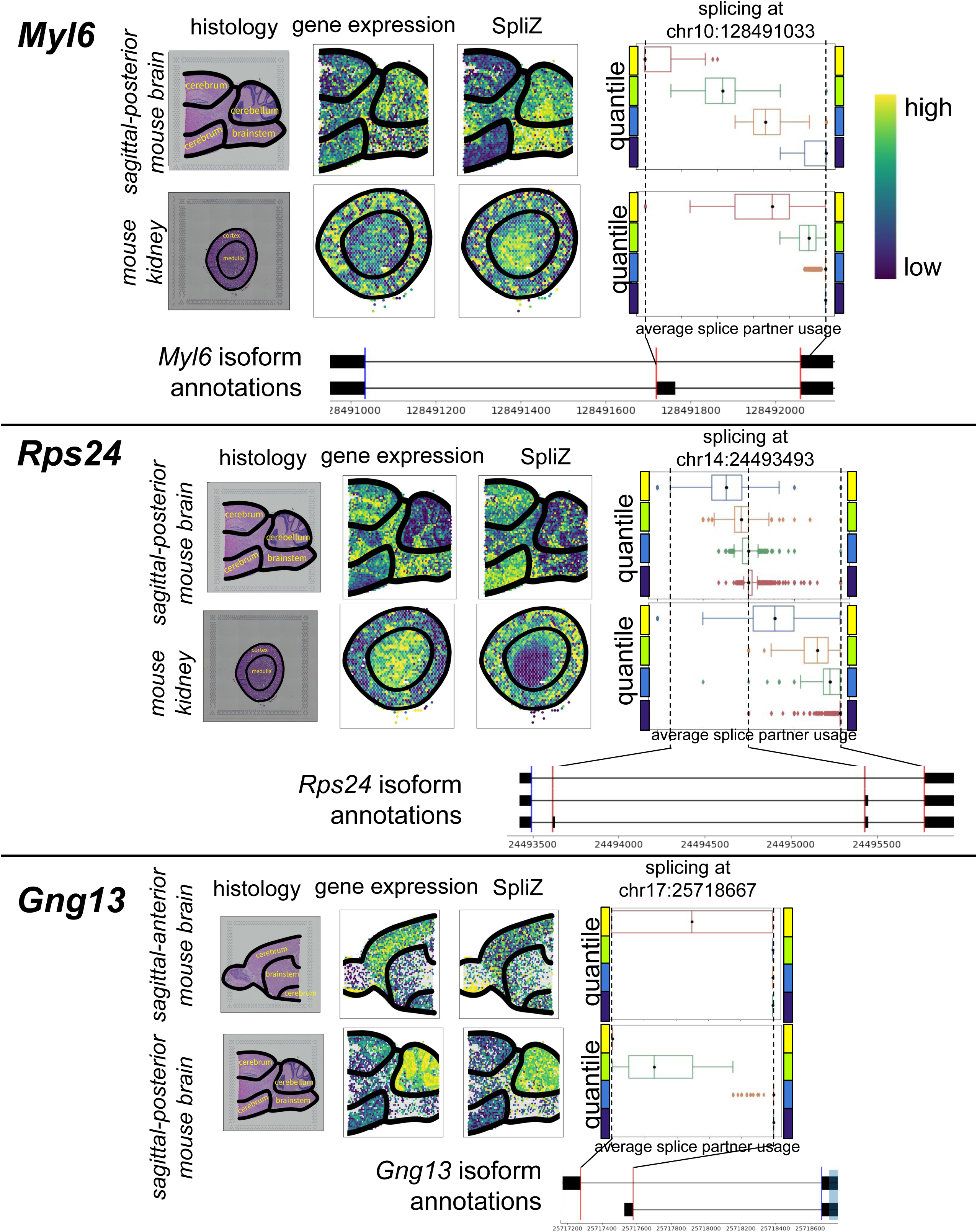
Splicing spatial regulation plots. The “histology” column contains histology images of the mouse sections, with anatomy annotated. In the “gene expression” column, each spot is colored according to its quartiled gene expression value. Dark blue corresponds to low gene expression, and yellow corresponds to high gene expression. In the “SpliZ” column each spot is colored by its quartiled SpliZ value. Box plots showing the splicing across each quartile are in the right-most column. For a single splice site (marked with a blue vertical line in the gene annotation), each Visium spot has some fractional usage of the corresponding splice sites (marked by red vertical lines in the gene annotation). Each box plot is created based on these fractions for the Visium spots in the given quartile. The whiskers on the box plots extend to the furthest point 1.5 times the interquartile range in either direction. All other points are marked as outliers. A) Splicing of *Myl6* is spatially regulated in the sagittal-posterior mouse brain and the mouse kidney. B) Splicing of *Rps24* is spatially regulated in the sagittal-posterior mouse brain and the mouse kidney. C) Splicing of *Gng13* is spatially regulated in the sagittal-anterior and sagittal-posterior mouse brain.

**Table 3:** Spatial score (Moran’s I) for each gene/window in each dataset. A) Scores for the SpliZ. B) Scores for the ReadZS. C) Scores for gene expression. D) Scores for gene expression by genomic window. Columns are: `gene` (gene name), `window` (genomic window identifier; use Table 4 to decode), `score_cont` (Moran’s I score), `num_pairs` (number of neighboring pairs with non-NA values for this gene/window in this dataset), `perm_pvals_emp_adj` (empirical permutation p value, adjusted by Benjamini Hochberg correction), `dataname` (name of the dataset).

The ribosomal protein gene *Rps24* was in the top three genes most significantly spatially regulated by Moran’s I for the SpliZ in all tissue sections (Figure 2A). Cell-type-specific splicing of *Rps24* is known to be ubiquitous in human, mouse and mouse lemur, including in the brain (Olivieri et al., 2021; Song et al., 2017). However, the spatial localization of *Rps24* splicing has not yet been shown. We observe higher levels of exon inclusion in the cerebrum of the mouse brain and the cortex of the kidney, compared to the rest of the tissues (Figure 3B; Figure 3—figure supplements 3 and 4).

The olfactory gene *Gng13* had one of the top-three highest Moran’s I for the SpliZ in both the sagittal-anterior and sagittal-posterior mouse brain (Figure 2A). *Gng13* is known to be highly expressed in the mouse cerebellum and outer layer of the olfactory bulb (Sanfilippo et al., 2021), but differential isoform expression has not previously been reported. We find that two different isoforms that differ by their 5’ end are expressed in the mouse brain, with the longer 5’ end mostly expressed in the tip of the cerebrum and the cerebellum, while the shorter 5’ end is mostly expressed in the rest of the brain (Figure 3C; Figure 3—figure supplements 5 and 6). In this case, lower levels of *Gng13* expression correlate with expression of the shorter isoform.

Two of the top three genomic windows with significant Moran’s I values for the ReadZS in the mouse brain were calcium-related genes with unannotated 3’ UTRs (Tables 3-4). The genomic window corresponding to the 3’ UTR of *Slc8a1* was within the top 3 most significant in the anterior brain, and had a larger Moran’s I value for ReadZS than gene expression in the sagittal-posterior mouse brain (Figure 2B). *Slc8a1* is a transmembrane protein that mediates the exchange of calcium and sodium ions. The mouse cerebrum contains mostly transcripts with an unannotated shorter 3’ UTR, while the cerebellum and brainstem mainly contain the annotated UTR (Figure 4A). Of interest, in some of these areas we see a third unannotated UTR that stretches beyond the annotated UTR (again spatially regulated, Figure 4A). Additionally, the calcium-channel-enabling gene *Gpm6a* was one of the top-two highest Moran’s I for the ReadZS in the mouse brain (Figure 2B). Gpm6a is a transmembrane protein present on neurons. It has been observed that overexpression of *Gpm6a* results in a Ca^2+^ influx, while *Gpm6a* inhibition leads to a suppression of Ca^2+^ influx and neuron differentiation (Michibata et al., 2008). In human hepatocellular carcinoma *GPM6A* has been shown to be regulated through binding of the miR-96-5p microRNA to the 3’ UTR of the gene, with production levels significantly decreasing with binding (Z.-R. Li et al., 2022). This indicates that differential UTR expression could result in fewer binding sites and thus affect *Gpm6a* production levels. The unannotated 3’ UTR is expressed more highly in the cerebrum of the mouse brain, while the annotated 3’ UTR is prevalent in the cerebellum (Figure 4B). Notably, high expression levels of *Gpm6a* are highly correlated with areas that include the shorter UTR, supporting the hypothesis that the shorter UTR doesn’t include the miR-96-5p binding site.

**Figure 4.**
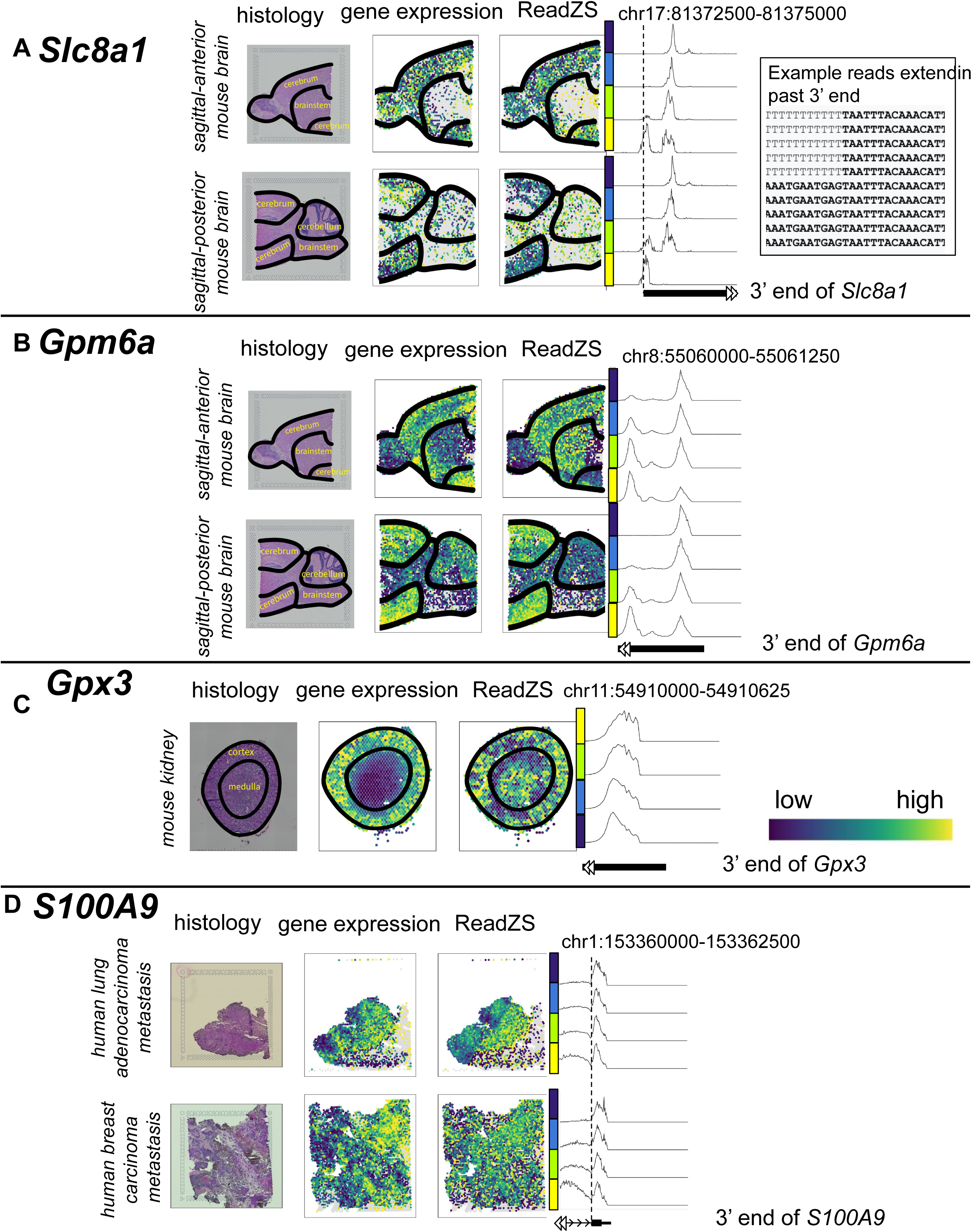
3’ UTR usage spatial regulation plots. The “histology” column contains histology images of the tissue sections, with anatomy annotated where possible. In the “gene expression” column, each spot is colored according to its quartiled gene expression value. Dark blue corresponds to low gene expression, and yellow corresponds to high gene expression. In the “ReadZS” column each spot is colored by its quartiled ReadZS value. Density plots to the right show the read density for the given genomic range, separated by ReadZS quantile. Color bars next to each peak plot indicate which quartile the distribution corresponds to. A) Unannotated 3’ UTRs of *Slc8a1* are spatially regulated in the sagittal-anterior and sagittal-posterior mouse brain. Several reads extend past the annotated 3’ end of the transcript. B) Unannotated 3’ UTRs of *Gpm61* are spatially regulated in the sagittal-anterior and sagittal-posterior mouse brain. C) Subtle differentiations in the 3’ UTR location of *Gpx3* are spatially regulated in the mouse kidney. D) Intron retention of *S100A9* is spatially regulated in human tumor brain metastases.

**Table 4:** Mapping of genomic windows to gene names. A) Mapping for mouse data. B) Mapping for human data. Columns are: `chr` (chromosome), `start` (beginning coordinate of the window), `end` (ending coordinate of the window), `window` (the name assigned to the genomic window), `strand` (the genomic strand), `gene` (the gene name(s) assigned to this window and strand).

*Gpx3* has the second-highest Moran’s I value in the mouse kidney for ReadZS (Figure 2B). Gpx3 is a selenoprotein secreted primarily by kidney proximal convoluted tubule cells. Transcripts at the interface of the cortex and the medulla have shorter UTRs compared to those in the cortex or the medulla. Only one 3’ UTR is annotated for *Gpx3*, but the 3’ UTR has been implicated in the rate of translation of *Gpx3* when selenium levels are low (BERMANO et al., 1996), and its 3’ UTR has been shown to confer stability (Wingler et al., 2001).

The RNA processing of housekeeping genes *Rps8* and *Actb* is also spatially regulated in the mouse brain. The ribosomal protein *Rps8* had the 6th-most-significant ReadZS spatial score in the sagittal posterior mouse brain. Although *Rps8* is not known to be differentially spliced, it has been shown that a diversity of ribosomal proteins helps create heterogenous selectivity of ribosomes (Shi et al., 2017). The distribution of reads among the exons of *Rps8* varies with spatial location (Figure 4—figure supplement 1), which could indicate different *Rps8* isoforms present. The actin gene *Actb* had the 8th-most-significant ReadZS Moran’s I score for the sagittal posterior mouse brain. Although Actin is considered a housekeeping gene, it is known to have an isoform with a longer UTR that confers higher translational efficiency of the transcript (Andreassi & Riccio, 2009; Ghosh et al., 2008). Our spatial analysis specifies the regions of the mouse brain and kidney that preferentially express the longer 3’ UTR, allowing functional understanding of which subtissues prioritize this transcript variant of *Actb* (Figure 4—figure supplement 2).

To show that we could identify spatial patterns in tissues without *a priori* anatomy annotations, we used this same methodology to identify spatial patterns in human tumor data (Sudmeier et al., 2021). We identified the calcium-binding protein *S100A9* as the 14th-most-significant ReadZS spatial pattern by Moran’s I in a brain metastasis from breast cancer carcinoma. The relative abundance of transcripts with a retained intron in *S100A9* varies spatially in both human lung adenocarcinoma and human breast carcinoma metastases (Figure 4D). *S100A9* is another calcium binding protein that is involved in the development of metastatic disease, though intron retention in this gene has not been previously described (Markowitz & Carson III, 2013). In this case the ReadZS identified an increase in intronic expression in some sections of the tumor sample. This serves as a taste of the rich discoveries that could be made by applying this methodology to unannotated tissue samples.

## Discussion

In addition to revealing previously unknown biology, this work serves as a proof-of-concept for analysis beyond differential gene expression localization from ST data. Future work exploring alternative methods of identifying spatially significant RNA processing patterns could likely identify more intricate spatial organization (for example, a method that would give a higher score to the SpliZ vs gene expression for *Gpx3*). Spatial analysis may be particularly revealing for the study of alternative splicing and alternate 3’ UTR usage due to the incomplete characterization of RNA isoforms: understanding where an isoform is expressed could inform hypotheses of isoform function. Because the 3’ UTR is known to contain motifs used to localize transcripts intracellularly, these methods will become all the more impactful when subcellular spatial sequencing data is available. Spatial analysis may be particularly fruitful when applied to tissue slices such as those from understudied organisms or cancerous tumors, because these datasets do not have previously-known anatomy. Models incorporating gene expression, splicing, and RNA processing information from multiple genes could help characterize these tissues in an annotation-free manner. Overall, this work represents a first step into the field of RNA processing analysis directly from ST data.

---

**Figure 2—figure supplement 1.**
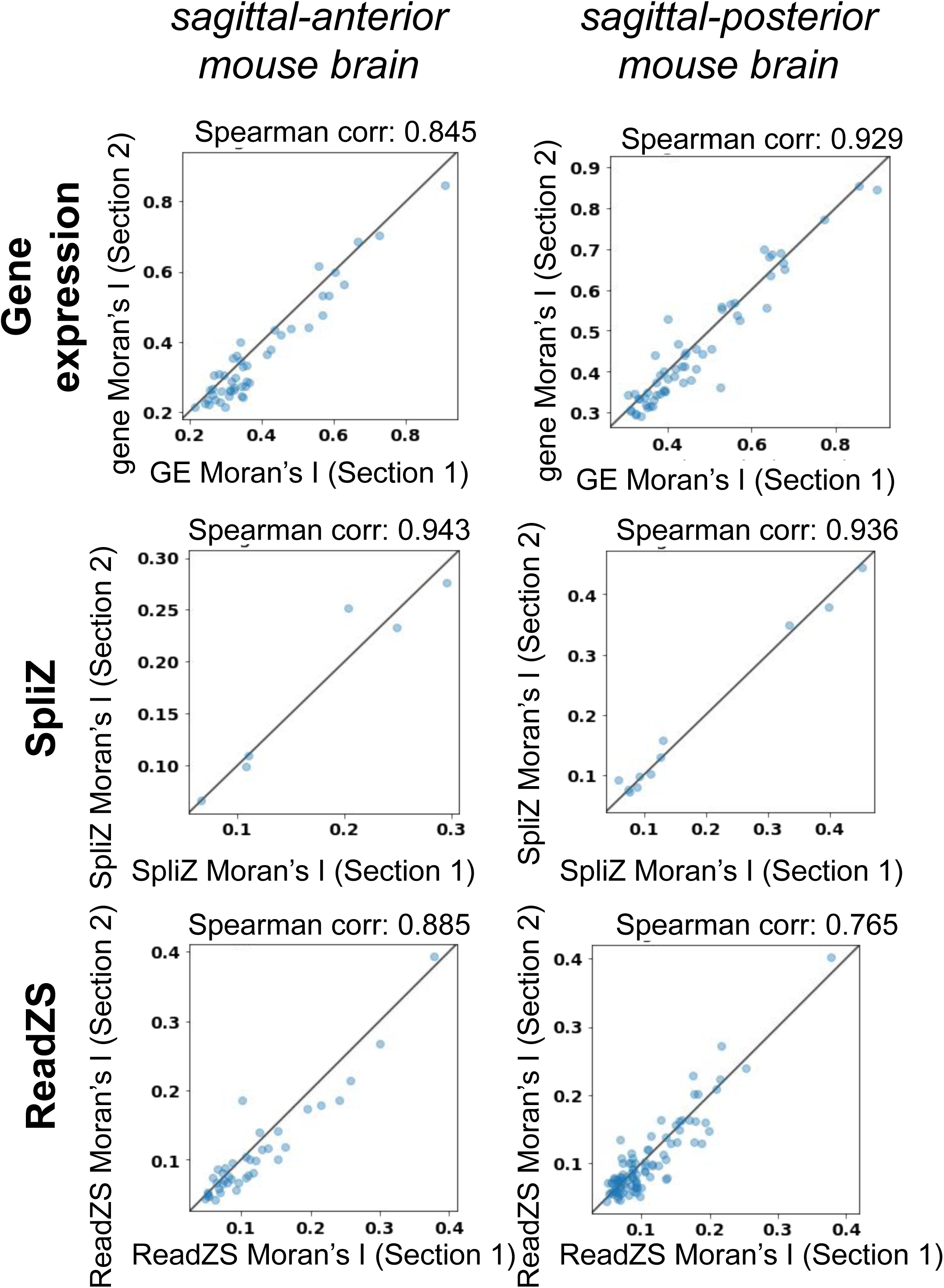
Moran’s I of biological replicates are highly correlated for gene expression, SpliZ, and ReadZS for those genes/genomic windows for which Moran’s I is significant in both replicates.

**Figure 2—figure supplement 2.**
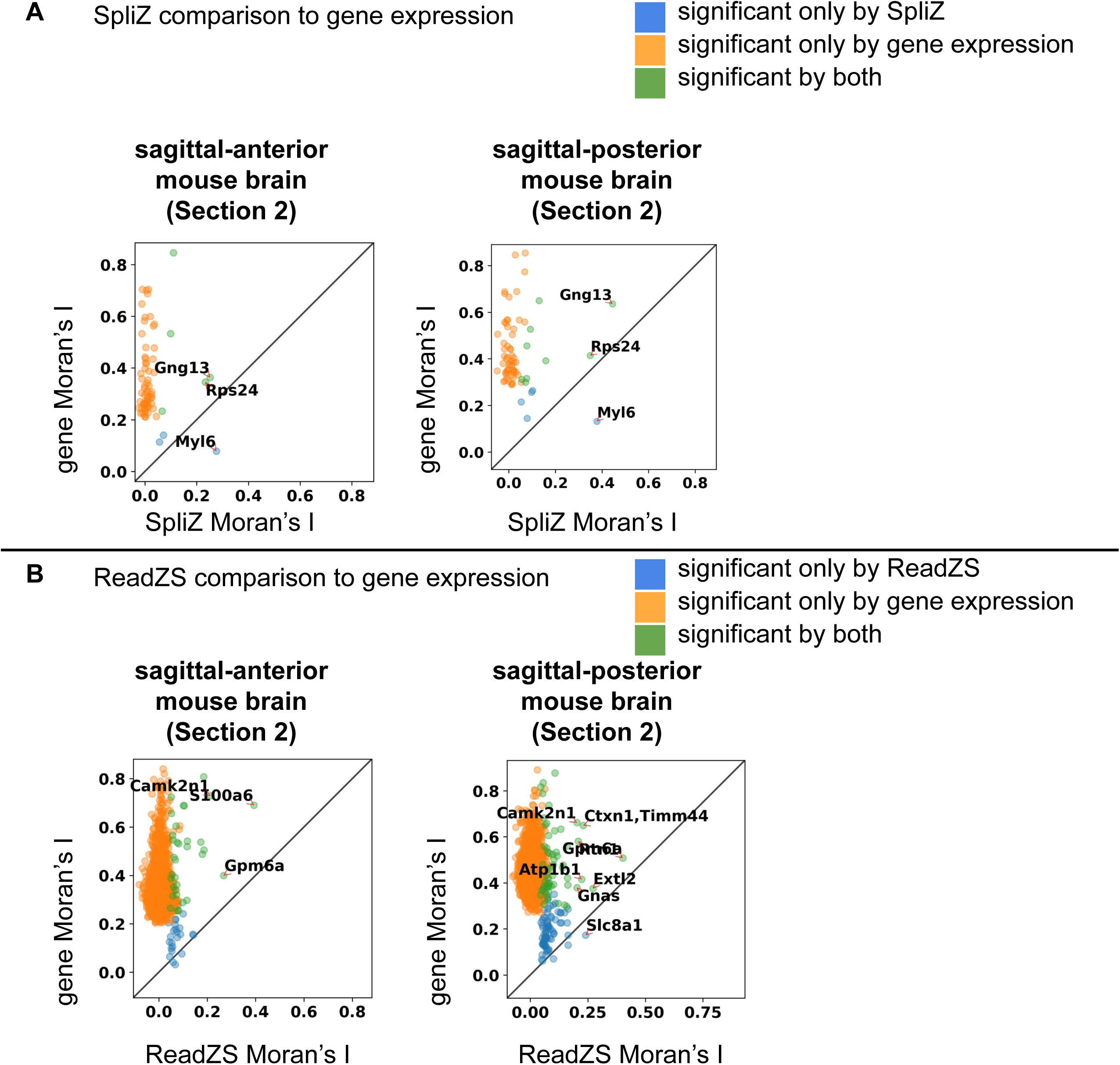
Biological replicates of Figure 2. A) Plots of SpliZ vs gene expression for sagittal-anterior mouse brain section 2 and sagittal-posterior mouse brain section 2. B) Plots of ReadZS vs gene expression for sagittal-anterior mouse brain section 2 and sagittal-posterior mouse brain section 2. See Figure 2 for a full description.

**Figure 3—figure supplement 1.**
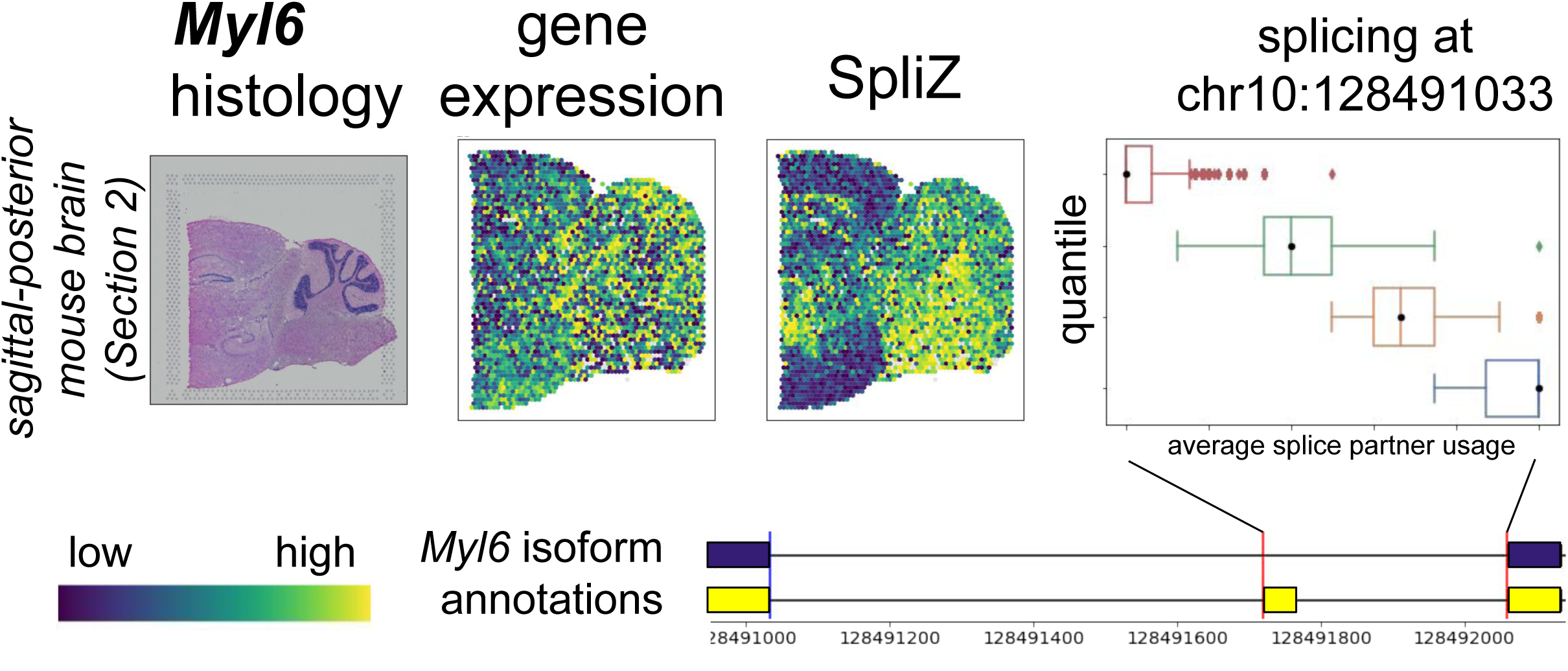
Biological replicate of *Myl6* splicing in the sagittal-posterior mouse brain (see Figure 3A).

**Figure 3—figure supplement 2.**
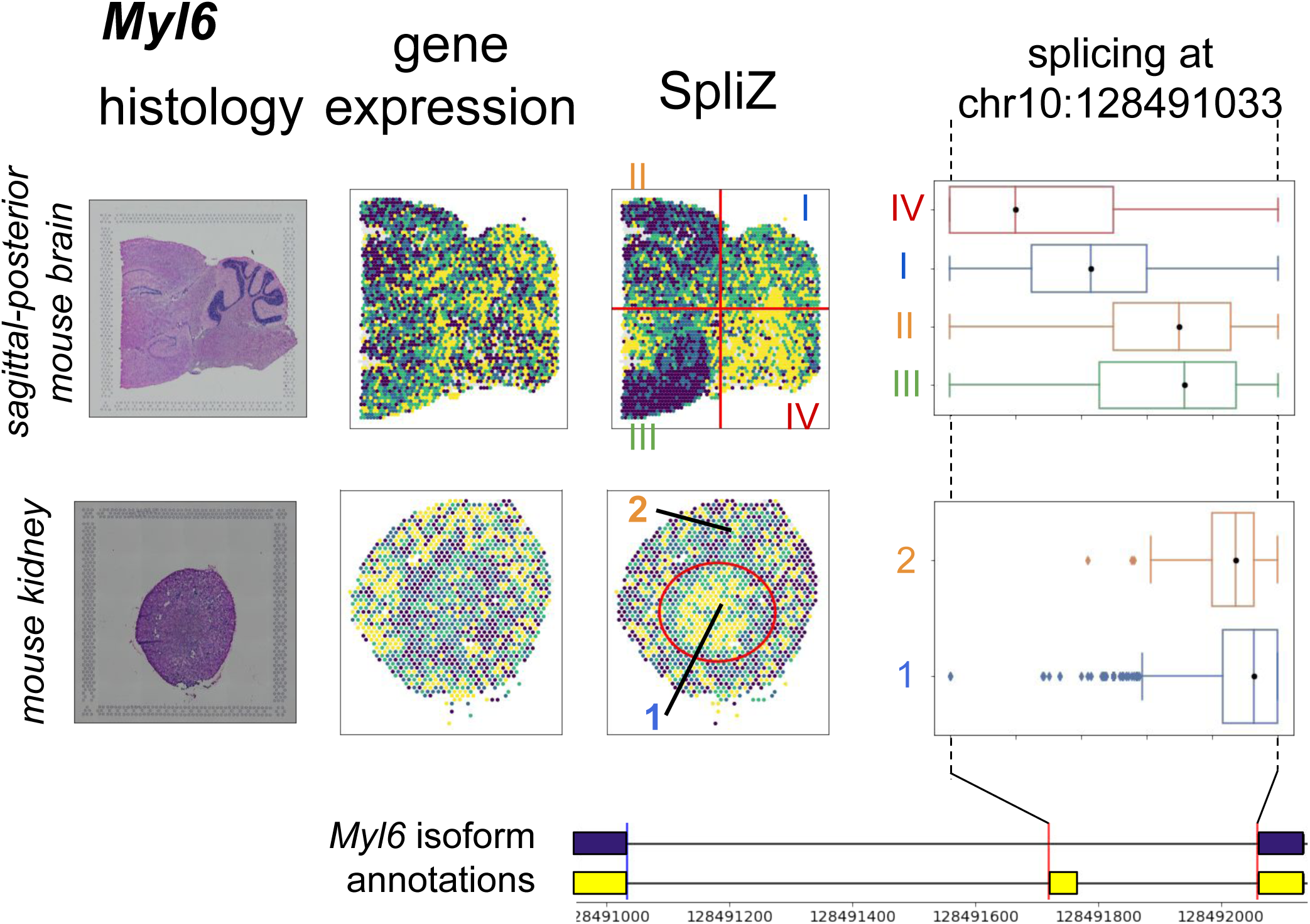
*Myl6* splicing in the sagittal-posterior mouse brain and mouse kidney, with box plots corresponding to the SpliZ values in each quadrant of the splicing plot and the inner vs outer cortex of the kidney, respectively.

**Figure 3—figure supplement 3.**
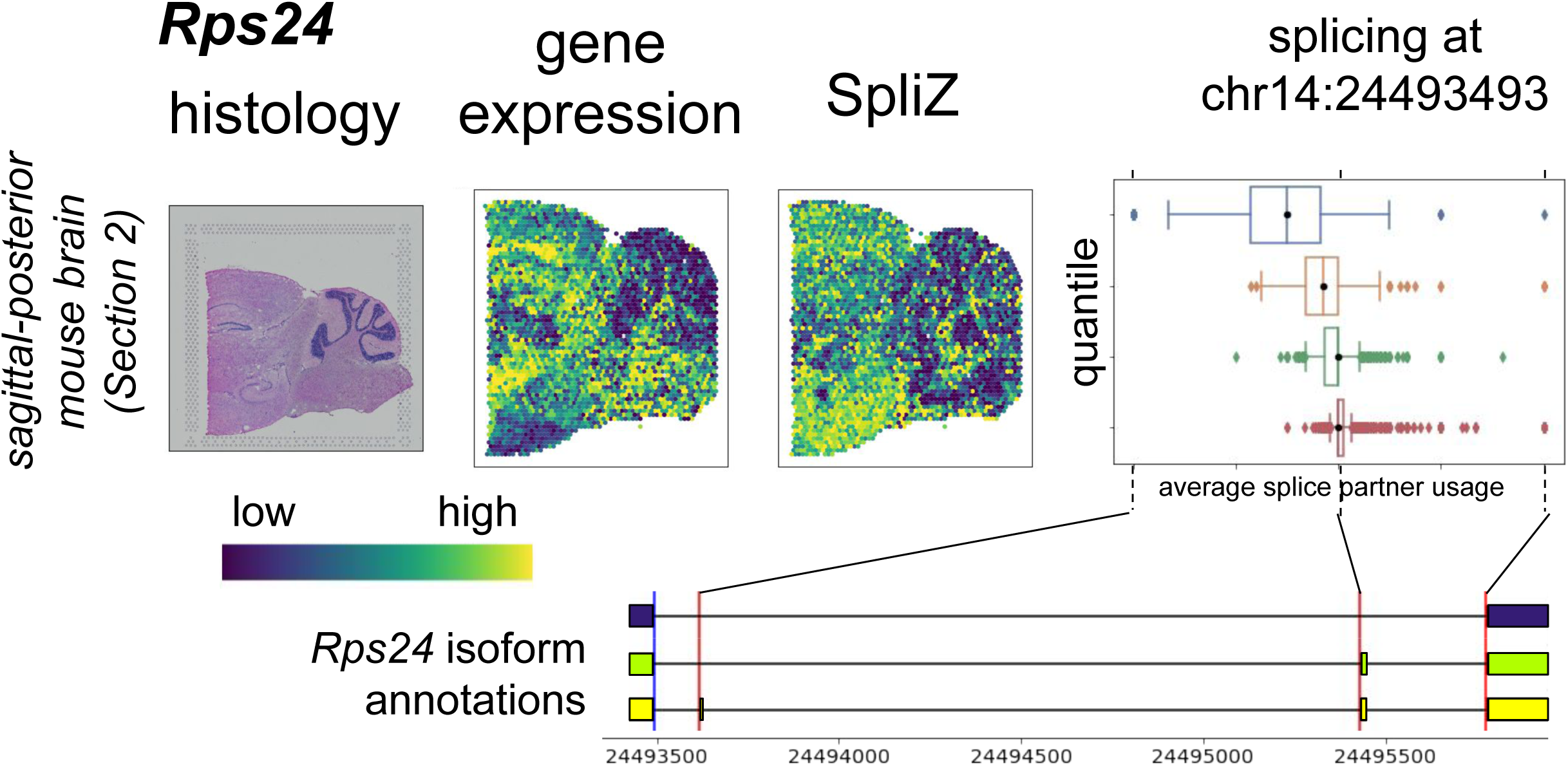
Biological replicate of *Rps24* splicing in the sagittal-posterior mouse brain (see Figure 3B).

**Figure 3—figure supplement 4.**
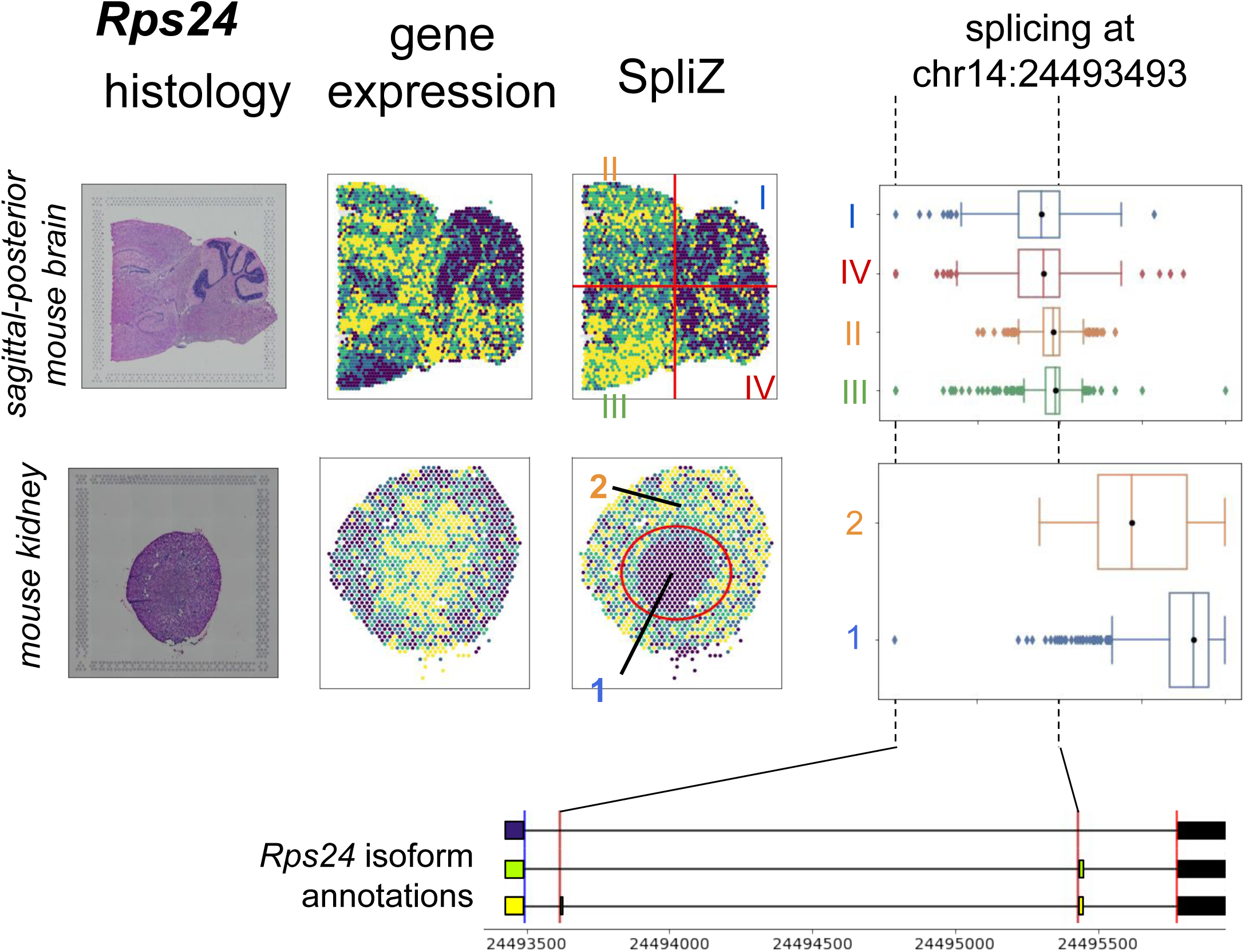
*Rps24* splicing in the sagittal-posterior mouse brain and mouse kidney, with box plots corresponding to the SpliZ values in each quadrant of the splicing plot and the inner vs outer cortex of the kidney, respectively.

**Figure 3—figure supplement 5.**
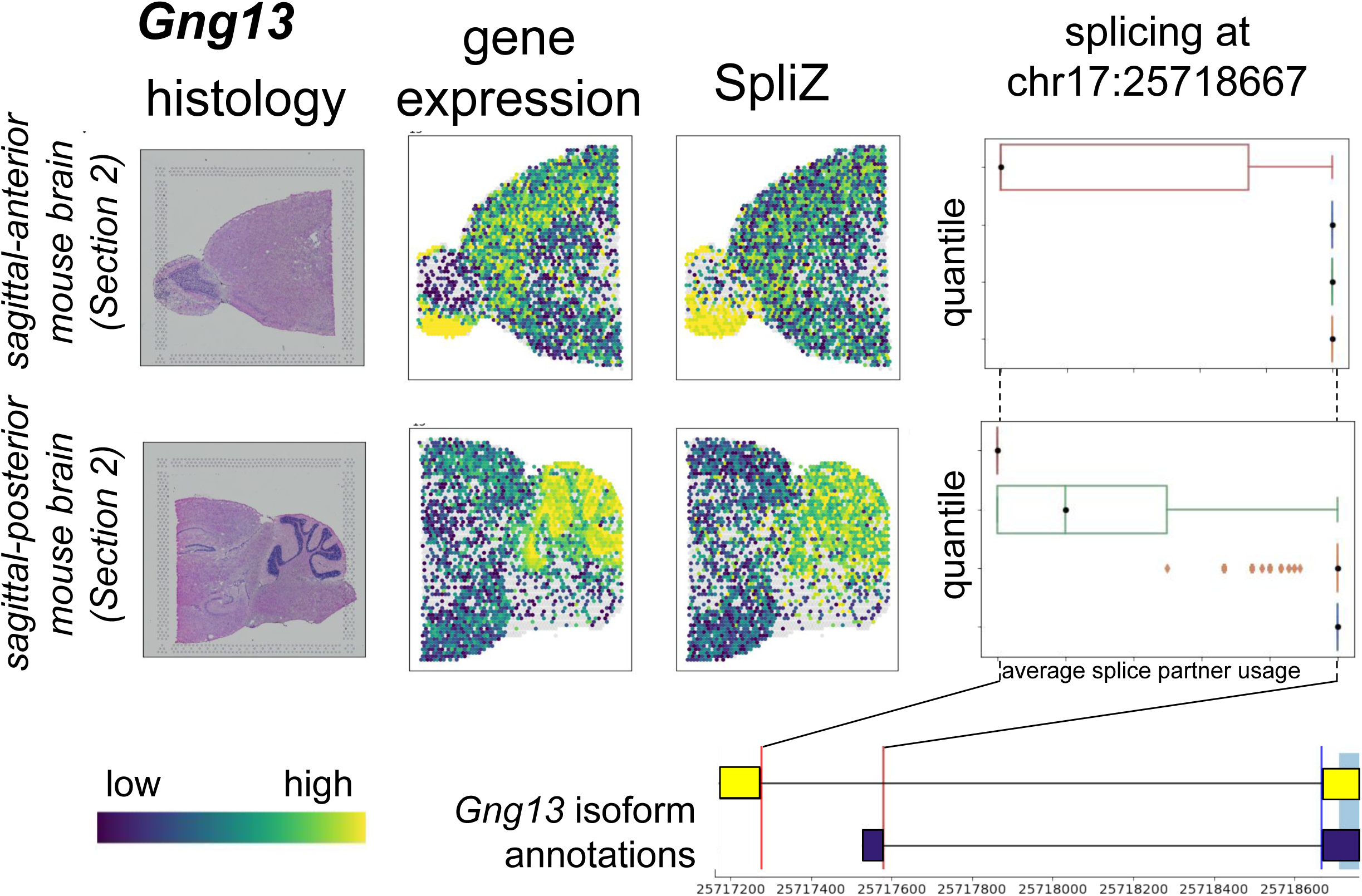
Biological replicate of *Gng13* splicing in the sagittal-anterior and sagittal-posterior mouse brain (see Figure 3C).

**Figure 3—figure supplement 6.**
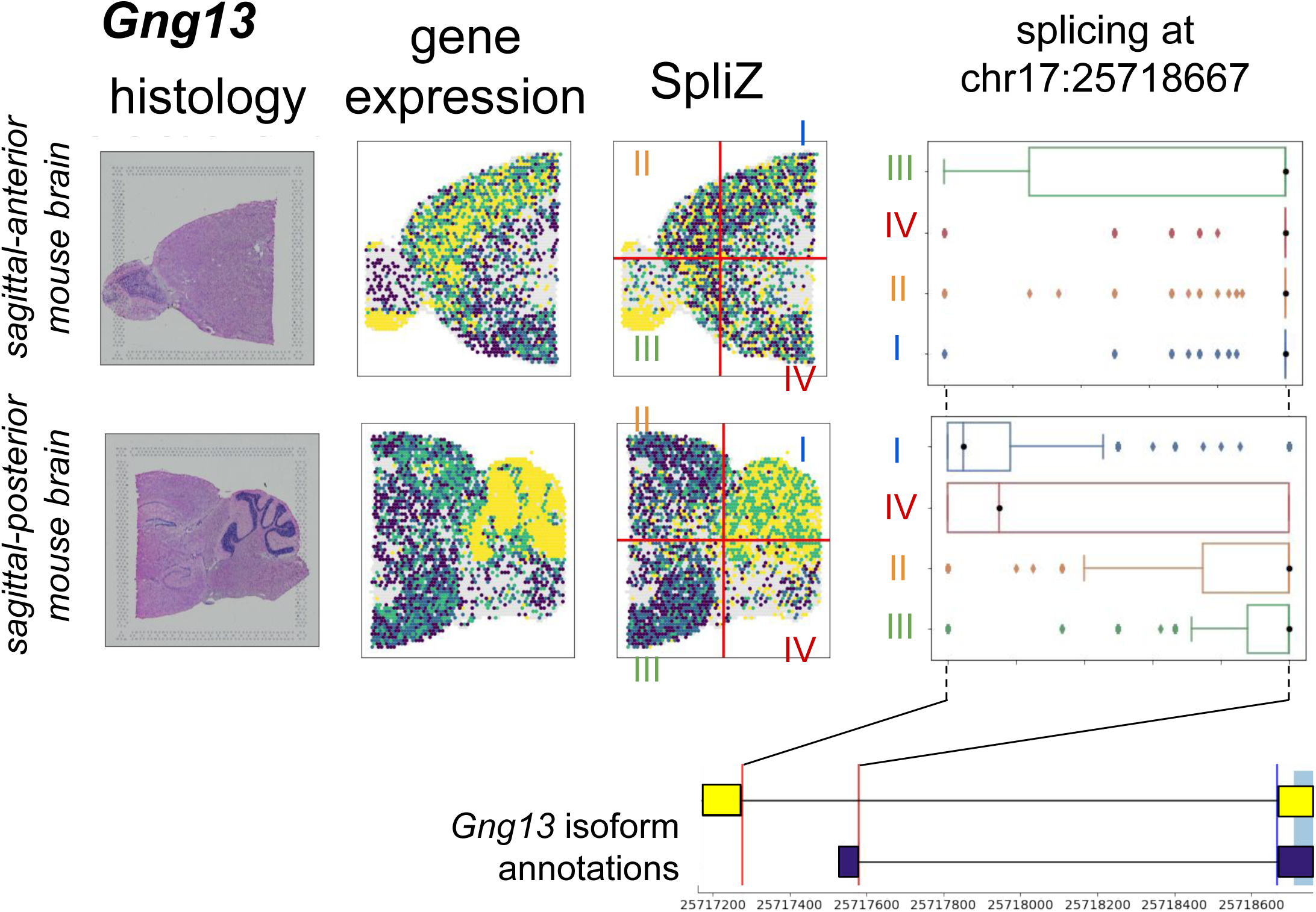
*Gng13* splicing in the sagittal-anterior and sagittal-posterior mouse brain, with box plots corresponding to the SpliZ values in each quadrant of the splicing plot for each.

**Figure 4—figure supplement 1.**
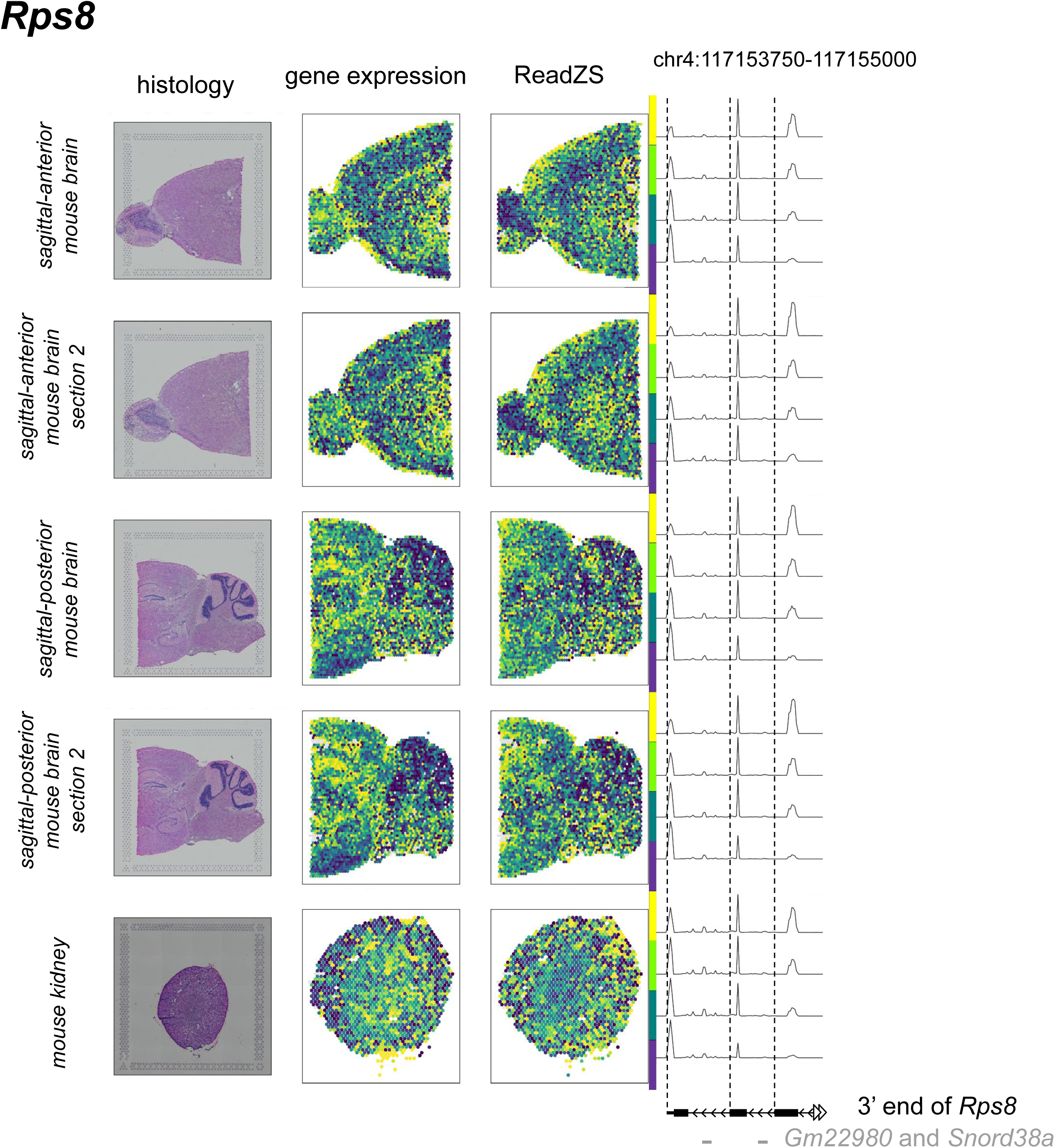
Exon expression of *Rps8* is spatially regulated in all four mouse brain sections and the mouse kidney. See Figure 4 for a full description.

**Figure 4—figure supplement 2.**
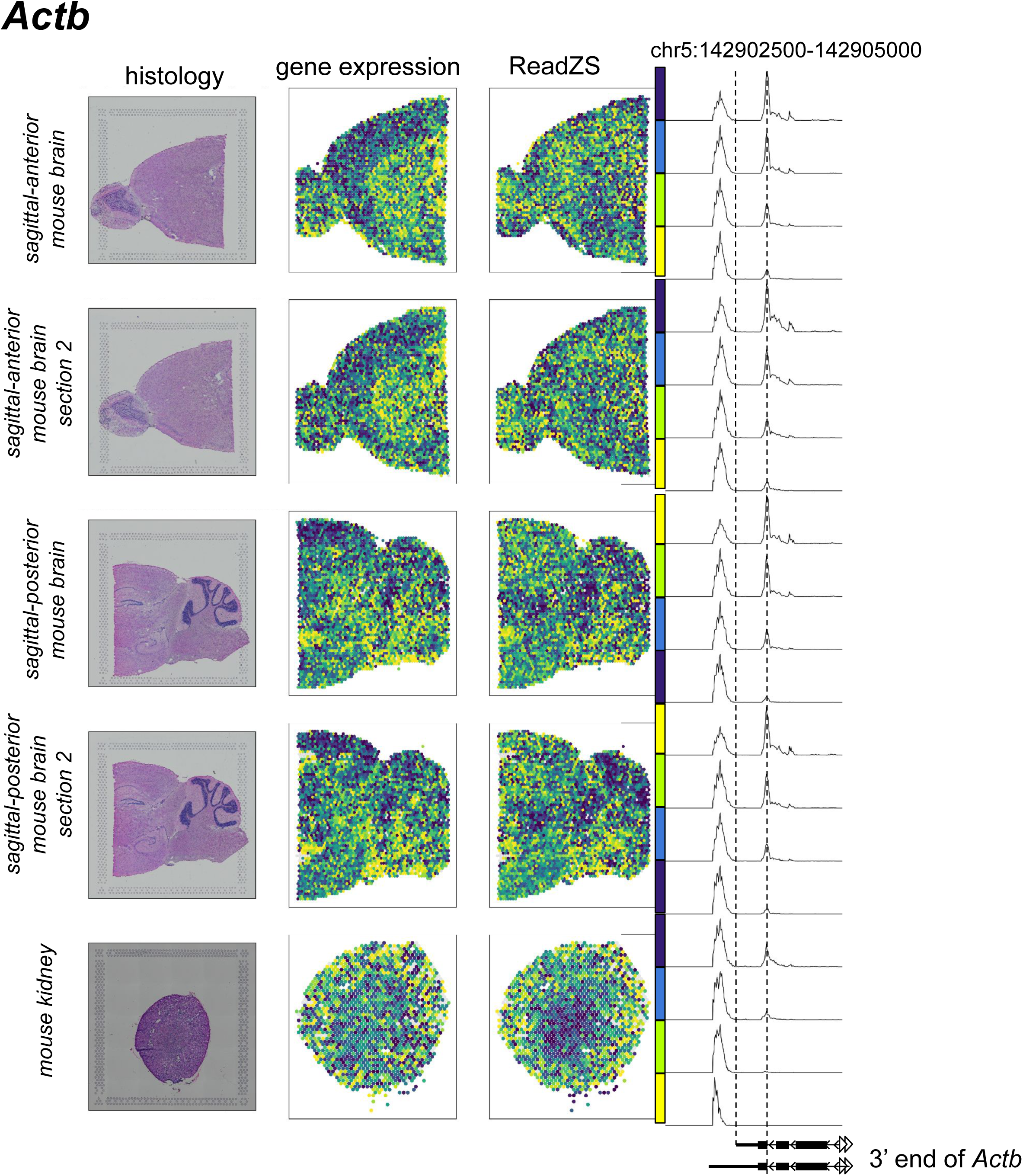
The 3’ UTR expression of *Actb* is spatially regulated in all four mouse brain sections and the mouse kidney. See Figure 4 for a full description.

## Table captions

All tables available here: https://doi.org/10.6084/m9.figshare.22144055.v1.

## Methods

## Acknowledgements

We thank the Salzman Lab for useful discussion. We thank Serena Y. Tan and Inma Cobos for helpful discussions surrounding the annotations of mouse and kidney histology images. JO is supported by the National Science Foundation Graduate Research Fellowship under Grant No. DGE-1656518 and a Stanford Graduate Fellowship. JS is supported by the Stanford University Discovery Innovation Award, National Institute of General Medical Sciences Grant R01 GM116847, and NSF Faculty Early Career Development Program Award MCB1552196. None of these funding sources were involved in study design, data collection and interpretation, or the decision to submit the work for publication.

## Data availability

BAM, gene expression, and spatial data was downloaded from the 10x genomics website for each dataset. This includes two sections of the sagittal anterior mouse brain (V1_Mouse_Brain_Sagittal_Anterior (sagittal-anterior mouse brain) and V1_Mouse_Brain_Sagittal_Anterior_Section_2 (sagittal-anterior mouse brain (Section 2))), two sections of the sagittal posterior mouse brain (V1_Mouse_Brain_Sagittal_Posterior (sagittal-posterior mouse brain) and V1_Mouse_Brain_Sagittal_Posterior_Section_2 (sagittal-posterior mouse brain (Section 2))), and one section of the mouse kidney (V1_Mouse_Kidney (mouse kidney)). A bash script to download all of the mouse data is available here: https://github.com/juliaolivieri/visium_analysis/blob/main/bash_scripts/download_mouse_data.sh. Human tumor data was downloaded from (Sudmeier et al., 2021) from GEO accession GSE179373. Data from p20218-s002_L2 (patient 27, lung adenocarcinoma) and p20218-s001_L1 (patient 26, breast carcinoma) was used in this study.

## Code availability

All custom code written for this paper is available here: https://github.com/juliaolivieri/visium_analysis.

## Running Spaceranger

BAMs were not included for the human tumor data, so Space Ranger v. 1.3.1 was used to create the BAMs. The human reference was downloaded using this command `curl -O` https://cf.10xgenomics.com/supp/spatial-exp/refdata-gex-GRCh38-2020-A.tar.gz. Space Ranger was run with default parameters. An example command is available here: https://github.com/juliaolivieri/visium_analysis/blob/main/scripts/submission_scripts/run_spaceranger.sh.

## Running the SpliZ

The SpliZ pipeline (https://github.com/salzman-lab/SpliZ) was run separately on each Visium slide with `bounds = 0` (requiring at least one read per gene per spot for the SpliZ score to be calculated), `grouping_level_1 = “dummy”`, and `grouping_level_2 = “pixquant`” (so differential splicing is determined based on quantiled pixel value of the image, though these differential splicing calls are not used), and otherwise default arguments (Olivieri et al., 2022). This treats each Visium spot as a “separate cell” and otherwise follows the logic of the original SpliZ paper (Olivieri et al., 2022). An example config file is available here: https://github.com/juliaolivieri/visium_analysis/blob/main/nextflow_inputs/visium_spliz.config, an example sample sheet is available here: https://github.com/juliaolivieri/visium_analysis/blob/main/nextflow_inputs/samplesheet_spliz.csv, and an example bash script is available here: https://github.com/juliaolivieri/visium_analysis/blob/main/nextflow_inputs/run_spliz.sh.

## Running the ReadZS

The ReadZS pipeline (https://github.com/salzman-lab/ReadZS) was run separately on each dataset with `ontologyCols = “pixquant”` (so differential RNA processing is determined based on quantiled pixel value of the image, though these differential RNA processing calls are not used), and otherwise default arguments (Meyer et al., 2022). This treats each Visium spot as a “separate cell” and otherwise follows the logic of the original ReadZS paper (Meyer et al., 2022). An example config file is available here: https://github.com/juliaolivieri/visium_analysis/blob/main/nextflow_inputs/visium_readzs.config, an example sample sheet is available here: https://github.com/juliaolivieri/visium_analysis/blob/main/nextflow_inputs/samplesheet_readzs.csv, and an example bash script is available here: https://github.com/juliaolivieri/visium_analysis/blob/main/nextflow_inputs/run_readzs.sh.

## Normalization of scores

We can create this “normalized” version of the SpliZ, ReadZS, or gene expression scores by doing the following for each gene/window:

1. Rank all values from smallest to largest (ties are broken randomly, so the same value can get multiple ranks)
2. Assign each rank to a uniform value between 0 and 1 (the value will be r/(R + 1) where *r* is the rank in question and *R* is the max rank of the dataset)
3. Use the reverse normal cdf to map each of these values to values from the normal distribution.

These “normalized” scores are used for analysis throughout the paper.

## Moran’s I value implementation

Moran’s I was calculated independently for each gene and each score as follows. Define a graph *G* for a given gene and score as follows: the vertices are spots with non-NA values for this gene/genomic window and score. There is an edge between two vertices if the corresponding pair of spots are next to each other (each internal spot is surrounded by six other spots). Let *s_i_* be the score (either SpliZ or ReadZS) of spot *i*. Then Moran’s I is defined by:

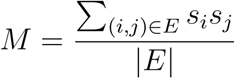

Because the SpliZ and ReadZS are zero-centered, under the null hypothesis that the score is not influenced by spatial location, the expected value of Moran’s I is 0. This value can also be calculated for gene expression values by performing the normalization procedure on the values first. The larger *M* is, the more spatial auto-correlation. Note that in this version we are not including influences of non-neighboring spots in the calculation.

## Permutations for Moran’s I

Permutation testing was used to assign p values to individual genes and scores. Note that “naive” permutation testing (permuting a single gene’s values and calculating *M* separately each time) is inappropriate, and causes the identification of many false positives. This is because we expect there to be some similarity between neighboring spots based on the experimental design, even under the null hypothesis: transcripts from the same cell can potentially be sequenced from two neighboring spots. We were instead interested in situations when this correlation between neighbors was unusually large.

We instead used the null where, for one permutation for a given gene, for each neighboring pair of spots with non-NA values for that gene, we randomly chose a gene for which both of those spots were non-null, and multiplied their scores.

We performed 1000 permutations with Benjamini Hochberg correction for each experiment. For analysis of the SpliZ, only genes with at least 100 non-NA values were used. For analysis of the ReadZS, only genomic windows with at least 1000 non-NA values were used. We used a threshold of 0.05 on the corrected p values to determine significance.

## Correlation between biological replicates

Correlation between biological replicates was assessed by comparing MBA1 to MBA2, and MBP1 to PMB2. For each pair, both discoveries were subset to those with adjusted p values less than 0.05. A spearman correlation was performed between the Moran’s I scores of the resulting genes/genomic windows.

## Analyzing gene expression to compare to the SpliZ

Gene expression values to compare with the SpliZ were extracted from the filtered gene expression matrices provided by Space Ranger. Gene expression scores were normalized in the same way as SpliZ and ReadZS scores, and spatial regulation was determined using the same method as well.

## Analyzing gene expression to compare to the ReadZS

Gene expression of genomic windows for comparison with the ReadZS was analyzed from the “counts” output of the ReadZS pipeline. Gene expression scores were normalized in the same way as SpliZ and ReadZS scores, and spatial regulation was determined using the same method as well.

## Correlation between gene expression and SpliZ/ReadZS on an individual gene/window level

For each given gene/window, a Spearman correlation was performed on the normalized SpliZ/ReadZS scores and the normalized gene expression scores for each spot.

## Annotation of mouse brain sections

Mouse brain sections were annotated through reference to the Allen Mouse Brain reference atlas (Sunkin et al., 2012, http://atlas.brain-map.org/atlas?atlas=2&plate=100883867#atlas=2&plate=100884125&resolution=14.44&x=7768.016736260776&y=4023.9998653017246&zoom=-3&structure=688&z=6). The mouse kidney was annotated with reference to the 3D virtual histology of murine kidneys (Missbach-Guentner et al., 2018).

